# Intraspecific variation in land use-related functional traits in *Plantago lanceolata*

**DOI:** 10.1101/2020.02.28.967521

**Authors:** Bence Gáspár, Oliver Bossdorf, Madalin Parepa

## Abstract

**Background and aims:** Intraspecific variation in functional traits is essential for the evolutionary success of organisms. The co-variation between trait variation and environment, as well as between different traits, can help us to understand which ecological factors drive habitat adaptation, and to what extent adaptation may be constrained by trait correlations and trade-offs. In managed grasslands, plants experience a combination of competition, recurrent biomass removal and nutrient pulses. Each of these ecological challenges requires specific plant tolerances, and populations should locally adapt if intraspecific variation exists in these traits.

**Methods:** We studied variation in land use-related traits in the common grassland plant *Plantago lanceolata*. In a common environment, we quantified the competitive ability (*R*\*), clipping tolerance and responses to a nitrogen pulse of plants from 54 populations with different land use intensities across Germany.

**Key results:** We found significant population differentiation in competitive ability but there was little evidence that trait variation was related to land use intensity. There was a positive relationship between competitive ability and clipping tolerance at the population level, indicating a genetic, and possibly functional, link between these two traits. In contrast, clipping tolerance and nitrogen responses were negatively correlated at the levels of plant individuals, indicating a physiological trade-off between plant responses to these two land-use processes.

**Conclusions:** Our results show that there is substantial intraspecific variation in some of the key functional traits for plant success in managed grasslands, and that rapid evolution and adaptation is therefore possible in these traits.

## Introduction

Understanding evolution in response to land use in grassland plants is of great interest because of the wide distribution and economic importance of these ecosystems, and because land use change is the strongest driver of global change (Foley *et al.*, 2005, Díaz *et al.*, 2019). Already from the early 20th century, grassland researchers showed that different management regimes resulted in rapid evolutionary changes in a range of grassland species. For instance, in a common-garden collection of over 400 *Dactylis glomerata* ecotypes, Stapledon (1928) found that there were persistent growth form differences between plants from different kinds of pastures and meadows. Later, Warwick and Briggs, in their classic studies on the “genecology of lawn weeds”, found similar results for several grassland species, e.g. dwarf, prostrate morphotypes originating from frequently mown lawns, and more erect ones in neighbouring populations that lacked the frequent mowing (Warwick & Briggs 1978, 1979). Evolution in response to land use was also found in the famous long-term Park Grass Experiment where Snaydon and Davies (1976) demonstrated local adaptation of *Antoxanthum odoratum* to different fertilisation and liming treatments (see also Davies & Snaydon 1973, 1976). In all of these classic studies, however, researchers compared simple categories of land use such as pastures versus meadows, or different types of fertilisation regimes, whereas finer-resolution analyses of land use processes are still rare. Moreover, previous studies usually focused on traits relevant for agriculture, such as yield, growth form and phenology, whereas other ecologically relevant functional traits received less attention.

From a plant eye’s view, three of the key processes in grasslands are (1) competition with neighbouring plants, (2) the temporary nutrient pulses created by animal droppings or fertilisation, and (3) the regular disturbance and biomass removal imposed by mowing or grazing. The abilities of plants to compete with neighbours, exploit nutrient pulses, and tolerate biomass removal are thus important functional traits in grassland plants.

Competition ultimately reduces the survival, growth or reproduction of an individual plant (Aarsen & Keogh, 2002), and plant species differ in the degree to which they they are impacted by neighbours (e.g. Keddy, 1990; Aarssen, 1992; Tokeshi, 2009). Plant competitive ability can be quantified in different ways (Aarssen & Keogh, 2002), and at the species level it appears to be particularly the ability to persist at low nutrient levels that makes some plant species outcompete others (resource ratio hypothesis; Tilman 1985). The significance of the so-called *R*\* value of species – the lowest resource level that allows persistence – has been proven by many species-level experimental studies (Wilson *et al.*, 2007). At the intraspecific level, a number of previous studies demonstrated genetically-based variation in competitive ability and the selective agency of neighbouring plants (Cheplick, 2015), but intraspecific variation in *R*\* has so far not been examined.

The second key process are nutrient pulses. Many ecosystems experience fluctuating resource availability, e.g. because of snowmelt, seasonal weather events or fires (Ostfeld & Keesing, 2000). Human activity is especially associated with such pulses, either indirectly through causing extreme climatic events (Coumou & Rahmstorf, 2012), or more directly through intentional nutrient deposition in agricultural landscapes. Resource pulses can impact population dynamics across communities and trophic networks (Gratton & Denno, 2003; Yang *et al.*, 2008) as well as across generations (Miao *et al.*, 1991), and they tend to promote particular plant species over others (Bilbrough & Caldwell, 1997), or even the spread of plant invaders (Parepa *et al.*, 2013). However, to our knowledge no previous study has investigated plant responses to nutrient pulses at the intraspecific level.

The third key process in grasslands is recurrent biomass removal. While strong mowing or grazing generally reduce fitness, plants possess the ability to regrow and to some extent compensate for such damage. Because of this, some species are able to maintain their fitness or even overcompensate and increase it in response to moderate levels of herbivory (McNaughton, 1983; Strauss & Agrawal, 1999). Plant tolerance to biomass damage has been extensively researched, and previous studies have repeatedly demonstrated not only species differences but also heritable variation within and among natural populations (e.g. Bergelson & Crawley, 1992; Agrawal, 1998; Strauss & Agrawal, 1999; Johnson, 2011), although rarely in relation to land use (but see Lennartsson *et al.*, 1997, 1998).

While all of the three described functional traits are expected to be important for success in managed grasslands, it seems unlikely that plants can evolutionarily improve all of them simultaneously. Increased competitive ability (= lower *R*\*) requires greater resource-efficiency, while stronger responses to nutrient pulses are only possible if plants are on the faster (= less resource-efficient) side of the fast-slow plant economy spectrum (Reich, 2014). Tolerance to biomass removal is usually based on belowground storage of resources, which means that some resources are not available for other purposes anymore. In general, we should expect evolutionary trade-offs (Agrawal *et al.*, 2010) between the three functional traits, and that the specific phenotypes evolving in different grasslands depend on the local intensities of fertilisation versus mowing and grazing damage.

We addressed these questions in the framework of the Biodiversity Exploratories project (www.biodiversity-exploratories.de), a large-scale and long-term network of ecological study sites for understanding relationships between land use, biodiversity and ecosystem functioning. The project includes 150 grassland plots across Germany (Fischer *et al.*, 2010), with 50 plots in each of the three regions Schorfheide-Chorin (northern Germany), Hainich-Dün (central Germany) and Schwäbische Alb (southwest Germany). Within each region, the plots cover a broad range of land use types and intensities. The detailed land use information available for these plots, with precise data on mowing frequencies, livestock densities and amounts of fertilisation, obtained from annual surveys (Blüthgen *et al.*, 2012), is a unique feature of the Biodiversity Exploratories project and, together with the large number of plots, makes it a powerful system for studying evolution in managed grasslands.

There is already evidence from the Biodiversity Exploratories that the phenotypes of several grassland species evolve in response to land use (Kloss *et al.*, 2011; Völler *et al.*, 2013, 2017). We built on these studies and examined 54 populations of the common perennial *Plantago lanceolata*. Unlike the previous studies, which only conducted simple phenotyping in a common environment, we carried out a greenhouse experiment with a series of treatments (Fig. 1) which allowed us to quantify the *R*\* values of plants, as well as their nutrient pulse responses and clipping tolerances. Specifically, we asked the following questions: (1) Is there intraspecific variation in the three functional traits in *P. lanceolata*? (2) What is the relationship between land use and the variation in these traits? (3) Are there trade-offs between the three traits, and are these trade-offs influenced by land use intensity?

**Figure 1.**
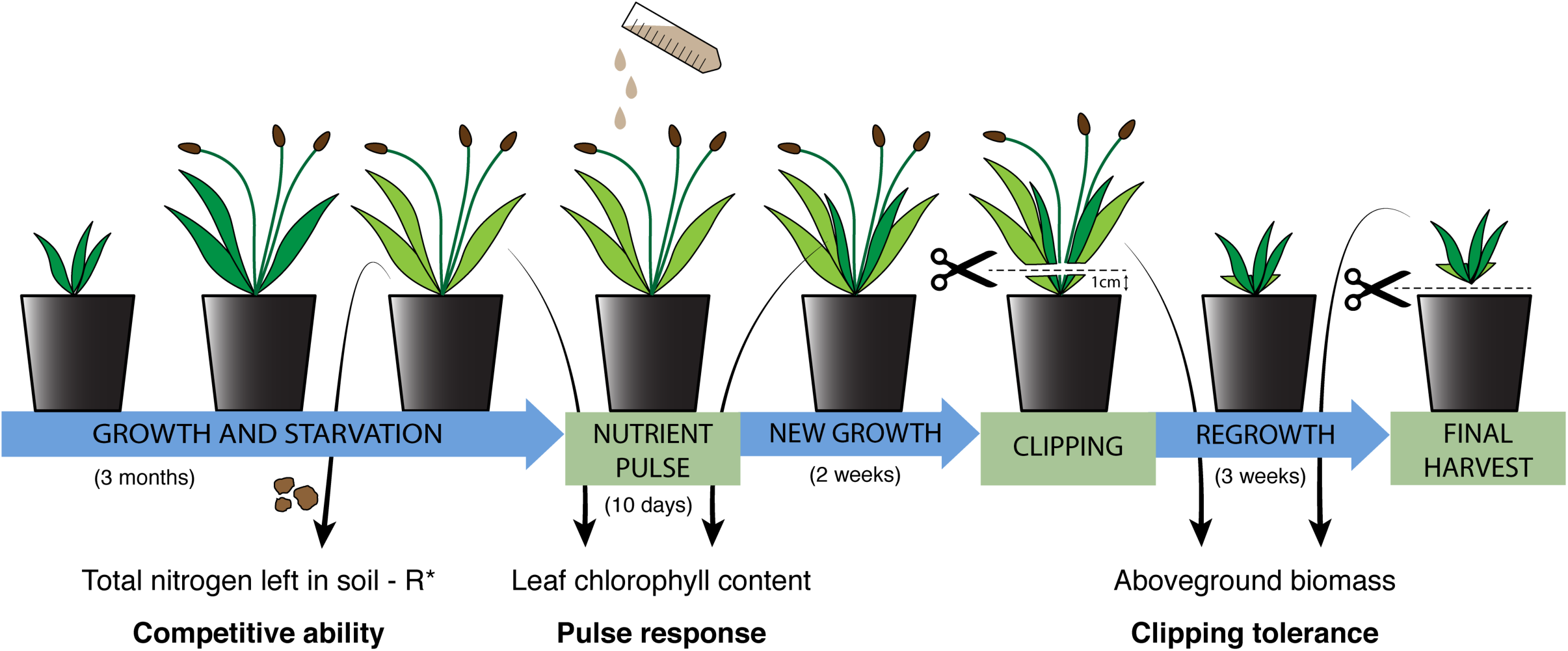
Schematic of the sequence and duration of experimental treatments used to estimate competitive ability (*R*\*), pulse response and clipping tolerance in *Plantago lanceolata* plants from 54 grasslands of different land-use intensities.

## Materials and Methods

### Study species and experimental design

To test the questions outlined above, we worked with *Plantago lanceolata* L. (Plantaginaceae), a short-lived perennial rosette herb that is very common in European grasslands and grows under a wide range of environmental conditions. *P. lanceolata* is also one of the most common plant species in the Biodiversity Exploratories, occurring on over 100 of the 150 grassland plots. In September 2015, we collected ripe seeds of *P. lanceolata* in each of the three regions, and from the broadest possible land-use gradient in each (Supplementary Table S2). Altogether, we sampled seeds from 54 plots, with 5–12 individual plants per plot.

We stratified the seeds at 5°C under moist and dark conditions for three weeks (Pons, 1992) and transplanted the germinated seedlings to 1-L pots filled with a 7:1.5:1 mixture of nutrient-poor soil, vermiculite and sand, with 5–12 individuals per population and a total of 540 plants (Supplementary Table S2). The pots were placed in a climate-controlled greenhouse with temperature set to 21°C/15°C at a 16h/8h day/night cycle. After six weeks, we rearranged all pots into a randomised block design, and we let the plants grow for another seven weeks to ensure strong nutrient depletion in all pots (Fig. 1). At this point, we took a 5 cm^3^ soil sample from each pot that was later analysed for total nitrogen content with a EuroEA Elemental Analyser (HEKAtech, Wegberg, Germany) at the Soil Biogeochemistry Lab at Karlsruhe Institute of Technology, and we measured the chlorophyll content of two leaves on each plant with a SPAD 502 chlorophyll meter (Konica-Minolta, Tokyo, Japan). After that, we fertilised each plant with 10 ml of liquid NPK fertiliser (Wuxal Universaldünger; Hauert MANNA Düngerwerke GmbH, Nürnberg, Germany) at a concentration equivalent to 50 kg N/ha. Ten days later, we measured chlorophyll content again on two newly grown leaves of each plant. Two weeks after adding the fertiliser, we clipped all plants one centimetre above ground. After another three weeks, we harvested the aboveground biomass of all plants, dried it at 70°C for three days, and weighed it.

### Data Analysis

Our data analyses generally focused on three variables: (1) the competitive ability of each plant, estimated as *1-R*\* (Tilman, 1985) where *R*\* was the fraction of total nitrogen in the potting soil left after 11 weeks of growth, (2) the pulse response as the ratio between the leaf chlorophyll contents after and before the fertilisation, with higher values indicating more successful utilisation of the added nitrogen, and (3) the clipping tolerance of plants, calculated as the ratio between their aboveground biomass from the second and first harvest, again with higher values indicating faster recovery from clipping damage.

Prior to the main analyses, we simplified our data by removing sources of variation that were not relevant to our study questions. We fitted linear models with the three regions of the Biodiversity Exploratories and the blocks in the greenhouse as fixed factors to each dependent variable, and we used the residuals from these models for all subsequent analyses (Manning *et al.*, 2015; Soliveres *et al.*, 2016). To improve the normality of error distributions, the data for pulse response and clipping tolerance were additionally log-transformed.

First, we tested for intraspecific variation in the three focus traits with mixed-effect models that included populations as fixed factors and maternal seed families nested within populations as random factors (Zuur *et al.*, 2009; see Supplementary Information Table S1 for model formulas). Second, we tested for relationships between land use and the three traits by fitting separate mixed models for each combination of land use intensities (mowing, fertilisation, grazing) and trait (competitive ability, pulse response, clipping tolerance), with each model including one of the land use intensities as explanatory variable plus population and maternal seed families nested within populations as random factors (Table S1). Next, we tested for trade-offs between the three focus traits by examining their statistical relationships at the level of individuals, seed families and populations. At the individual-level, we fitted mixed models with random intercept and slope that included the respective other trait as explanatory variable, plus population and family nested within population as random factors. At the family level, we analysed family means and included only population as random factor, and at the population level, we used simple linear models regressing the population means of two traits against each other. In the cases where we found significant relationships between the traits, we proceeded to the final step in our analyses where we tested the effects of land use on trait relationships. We did this through a series of mixed models with random intercepts and slopes that included the respective other trait, one of the three land use intensities, and their interactions, as fixed factors, plus populations and families nested within populations as random factors. All statistical analyses were done in R (R Development Core Team, 2008). We corrected all *P*-values for false discovery rates (FDR).

## Results

We found significant heritable variation, both at the population and seed family level, for competitive ability, but only marginally significant family-level variation in clipping tolerance, and no significant variation at all in pulse response (Table 1 and Figure 2). There were no significant relationships between land use intensity and the three studied functional traits (Table 2). When we tested for relationships between competitive ability, pulse response and clipping tolerance, we found significant negative relationships between pulse response and clipping tolerance at the level of individuals and seed families, and a significant positive relationship between competitive ability and clipping tolerance at the population level (Table 3 and Fig 3). Furthermore, we found a significant effect of mowing on the individual-level relationship between pulse response and clipping tolerance (*F* = 9.08, *P* = 0.025 for mowing x pulse response interaction), with the negative relationship between the two traits disappearing at higher mowing intensities (Fig 4). There were no other significant land use effects on trait relationships.

**Table 1.** Results of mixed models testing for heritable variation in three functional traits in *Plantago lanceolata*, with populations and seed families as fixed and random factors, respectively. LRT = likelihood-ratio test. All *P*-values are FDR-corrected.

|  | Population |  | Seed family |  |
| --- | --- | --- | --- | --- |
|  | *F* | *P* | *LRT* | *P* |
| Competitive ability | 1.60 | **0.049** | 16.82 | **0.000** |
| Pulse response | 1.01 | 0.463 | 0.04 | 0.835 |
| Clipping tolerance | 1.30 | 0.169 | 4.05 | 0.066 |

**Table 2.** Results of mixed models testing for the effects of mowing, fertilisation and grazing on three functional traits in *Plantago lanceolata*, with populations and seed families included as random factors. Est. = slope estimate of the models. All *P*-values are FDR-corrected.

|  | Mowing |  |  | Fertilisation |  |  | Grazing |  |  |
| --- | --- | --- | --- | --- | --- | --- | --- | --- | --- |
|  | Est. | *F* | *P* | Est. | *F* | *P* | Est. | *F* | *P* |
| Competitive ability | 0.142 | 4.31 | 0.387 | 0.050 | 1.51 | 0.504 | -0.071 | 1.68 | 0.504 |
| Pulse response | 0.014 | 0.08 | 0.838 | 0.001 | 0.04 | 0.838 | 0.009 | 0.06 | 0.838 |
| Clipping tolerance | 0.007 | 1.85 | 0.838 | 0.011 | 1.60 | 0.504 | 0.007 | 0.39 | 0.838 |

**Table 3.** Results of random slope and intercept mixed effects models testing for relationships between competitive ability (CA), pulse response (PR) and clipping tolerance (CT) in *Plantago lanceolata*, with populations and seed families included as random factors. Est. = slope estimate of the models. All *P*-values are FDR-corrected, and *P*<0.05 are in bold.

|  | Individuals |  |  | Families |  |  | Populations |  |  |
| --- | --- | --- | --- | --- | --- | --- | --- | --- | --- |
|  | Est. | *F* | *P* | Est. | *F* | *P* | Est. | *F* | *P* |
| CA ~ PR | 0.07 | 3.26 | 0.162 | 0.12 | 1.31 | 0.336 | 0.01 | 0.00 | 0.967 |
| CA ~ CT | -0.01 | 0.01 | 0.967 | 0.11 | 1.33 | 0.336 | 0.44 | 6.21 | **0.048** |
| PR ~ CT | -0.05 | 11.47 | **0.009** | -0.05 | 8.81 | **0.018** | -0.05 | 2.52 | 0.212 |

**Figure 2.**
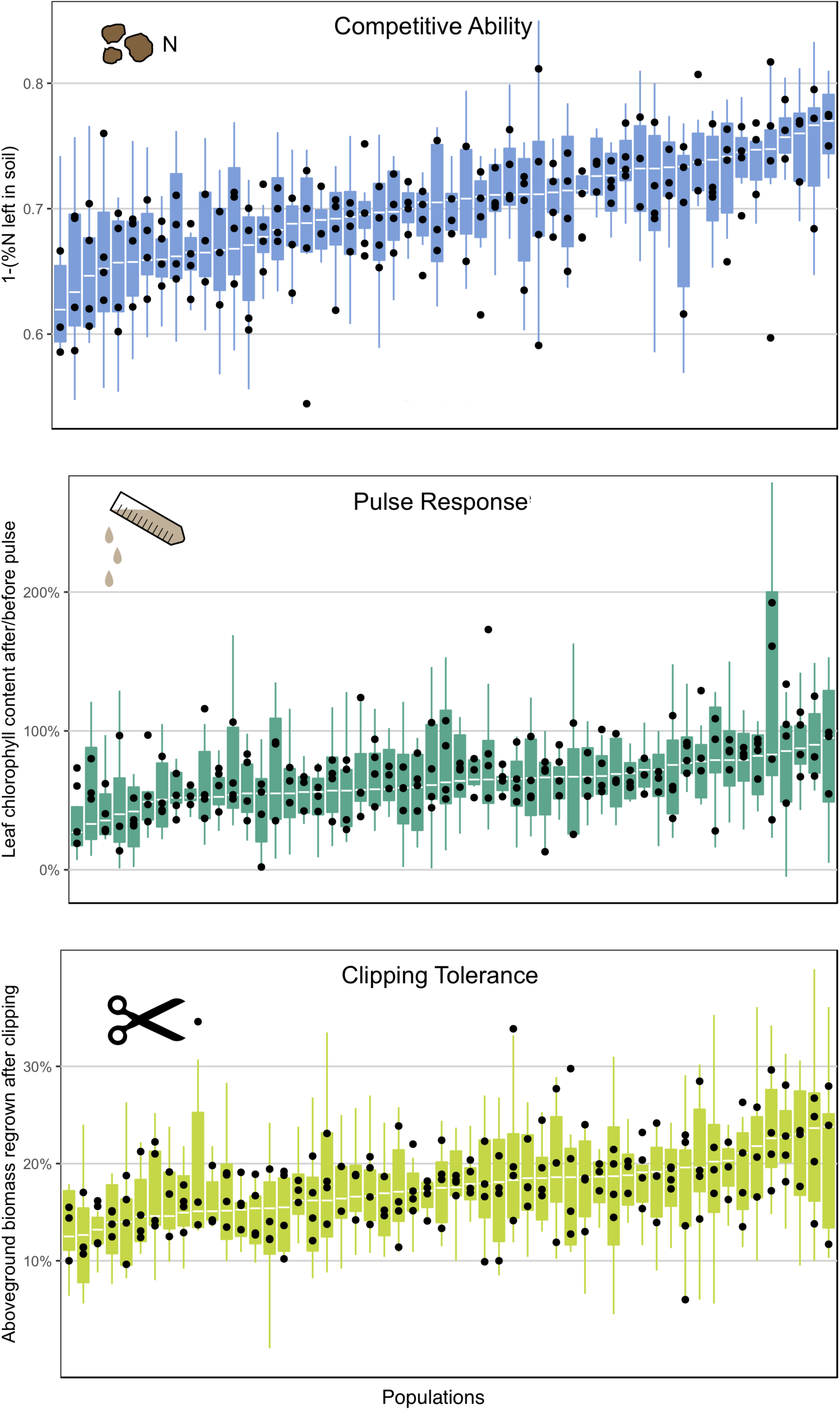
Variation among populations (boxplots) and seed families (black dots within boxplots) in three functional traits in *Plantago lanceolata*. The boxplots are based on all individuals per population and indicate medians, 25th/75th percentiles, and the 1.5 x interquartile ranges. For each trait, populations are ordered by their median values.

**Figure 3.**
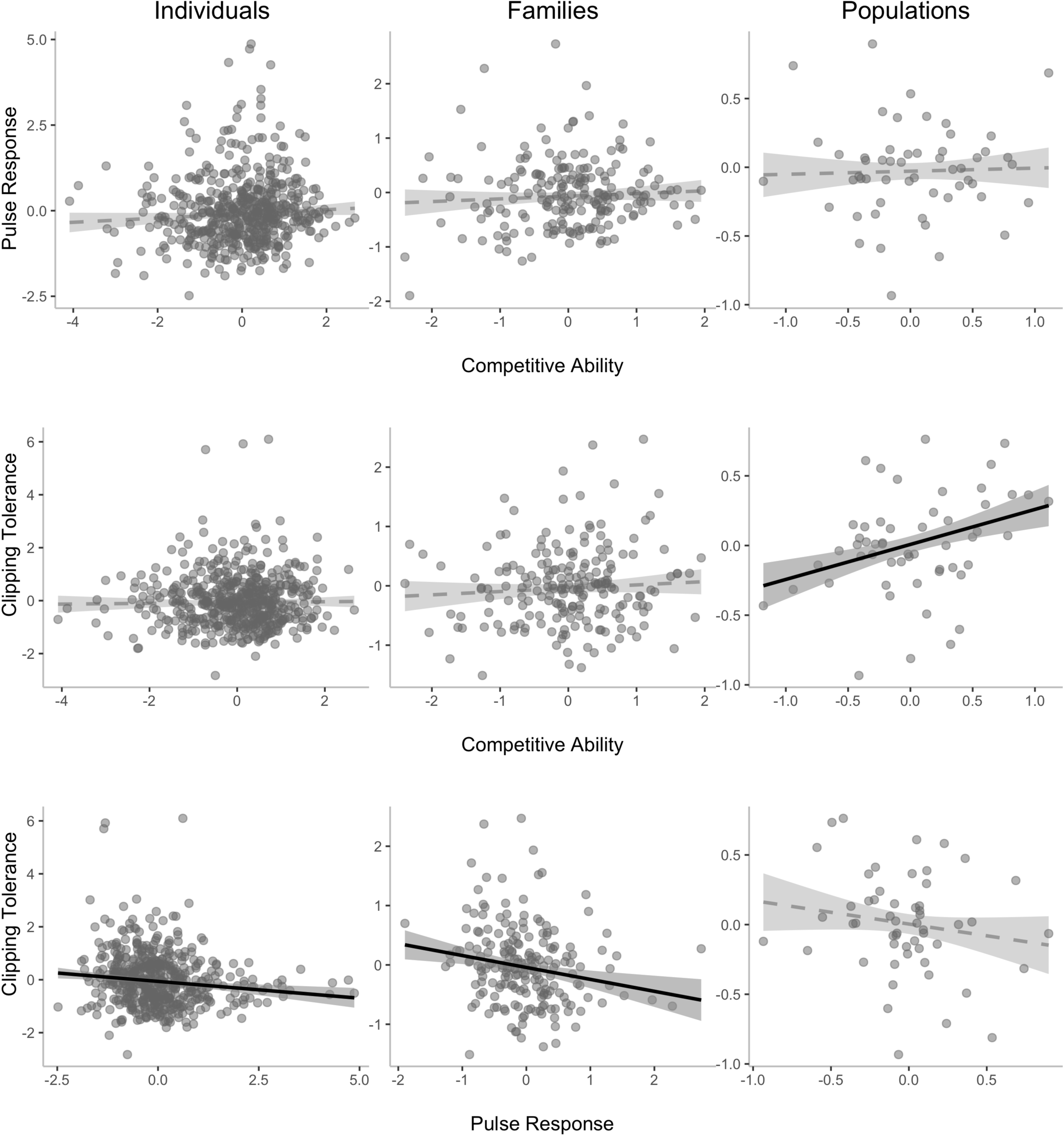
Relationships between the three studied functional traits of *Plantago lanceolata* at the levels of individuals, seed families and populations. Solid and dashed line plots indicate the fitted models for significant and non-significant relationships, respectively, with their 95% confidence intervals.

**Figure 4.**
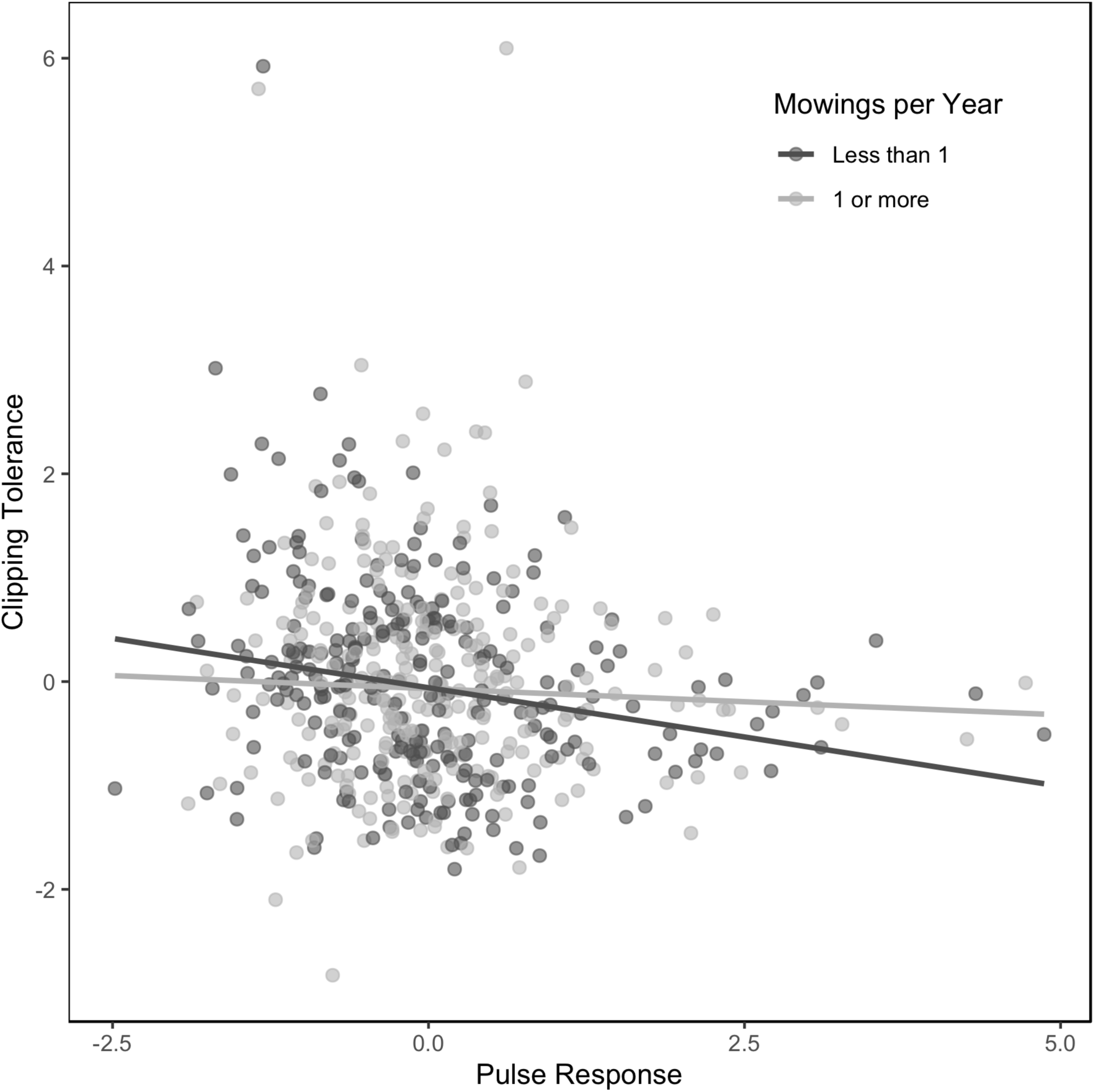
The mowing intensity of their grasslands of origin affects functional trait correlations in *Plantago lanceolata*. Each dot represents a plant individual grown in a common environment.

## Discussion

To understand plant intraspecific variation in relation to land use, we studied 54 grassland populations of *Plantago lanceolata* that strongly differed in their intensities of mowing, grazing and fertilisation. We specifically examined three functional traits that we expected to be important for plant survival in grasslands: competitive ability, clipping tolerance and the ability of plants to quickly respond to nutrient pulses. We found substantial intraspecific variation in competitive ability (*R*\*) but not in the other two traits, and there was no evidence for population-level relationships between traits and land-use intensity. However, there were several positive or negative relationships between functional traits at the levels of individuals, families or populations, indicating physiological or evolutionary links between these traits. Below, we discuss the results in detail, and attempt to place them into a broader context.

### Intraspecific variation

A necessary prerequisite for genetic differentiation and local adaptation in the examined traits is that our study system harbours significant intraspecific variation. We did not find any significant family- or population-level variation in clipping tolerance and plant responses to a nutrient pulse, but there was substantial intraspecific variation in *R*\* competitive ability, both at the level of seed families and populations. To our knowledge, this is the first time that intraspecific variation, and thus microevolution, in this aspect of competitive ability has been studied and demonstrated in plants.

We were surprised to not find population differentiation in clipping tolerance because intraspecific variation has been repeatedly shown in other plant species (e.g. Agrawal *et al.*, 1999; Johnson, 2011; Juenger & Bergelson, 2000; Strauss & Agrawal, 1999; Deng *et al.* unpublished). We also found no population differentiation in pulse response, and there are no previous studies on intraspecific variation in this trait. With 54 populations and 199 seed families, a lack of statistical power is an unlikely explanation in our case. Instead, we think that it may have been a combination of weak true patterns and high signal-to-noise ratio. First, since *Plantago lanceolata* is wind-pollinated and self-incompatible (Kuiper & Bos, 1992), there is generally strong gene flow and relatively weak population differentiation in this species (Gáspár *et al.* 2019). Second, we worked with an F_1_ generation that had random fathers (from the field) but that, unlike under field conditions, was not experiencing strong natural selection. This likely further increased variation among individuals and therefore lowered the signal-to-noise ratio in our system. Finally, clipping tolerance and pulse response are both derived traits based on several, error-prone measurements, and thus error propagation could have further added to this problem. However, in spite of all this, we did find significant family- and population-level variation in *R*\*, which underlines the ecological and evolutionary significance of this result.

### No relationships with land use

We found no relationships between the land-use intensities recorded in the Biodiversity Exploratories and the three studied functional traits. This contrasts with previous studies in the Biodiversity Exploratories (Völler *et al.*, 2013, 2017) as well as in other systems that demonstrated land use-related phenotypic changes in plants (e.g. Aarssen & Turkington, 1985a, b, c, 1987; Lennartsson *et al.*, 1997; Briggs, 2009). In principle, there are three possible explanations: (1) a true pattern could not be detected because of statistical or methodological shortcomings, (2) there was no pattern yet because the land use has not been acting long enough yet in our system, or (3) there is no pattern. As already explained above, our study did not lack statistical power, and it covered a broad range of land use intensities, also compared to previous studies. Moreover, although there is some interannual variation in land use in the Biodiversity Exploratories (Blüthgen *et al.*, 2012; Allan *et al.*, 2014), which could potentially impede the impacts of natural selection, previous studies already found land use-related differentiation of plant phenotypes in our system (Völler *et al.*, 2013, 2017). It is also known from other studies that that a couple of years can be enough for stable shifts in plant phenotypes between differential management (Briggs, 2009). Therefore, explanations (1) and (2) both appear unlikely, and we need to consider the third option that there might simply be no relationships between land use and the three studied functional traits; possibly because of the derived nature of the traits, or evolutionary constraints particular to these traits and land use in this system.

### Correlations between three functional traits

Besides quantifying intraspecific variation in the three functional traits and their relationships with land use, we also tested for interrelationships between traits, and we did this at three levels: plant individuals, maternal seed families and populations. Each of these levels provides us with different answers: at the level of individuals, trait correlations are most likely related to functional-physiological constraints or necessities, whereas at the level of seed families they reflect underlying genetic correlations, and at the level of populations they rather indicate trait syndromes associated with habitat adaptation.

We found no relationships between competitive ability and pulse response at any of these levels. This was surprising as lower *R*\* values (i.e. better competitive ability) should be coupled to a resource-conservative plant economy, whereas strong responses to nutrient pulses require a large metabolic capacity. We therefore expected a trade-off between the two traits. However, our results suggest that competitive ability evolves independently. The only observed trait correlation involving competitive ability was a positive population-level correlation between competitive ability and clipping tolerance, indicating both traits might be beneficial in the same environments. Resprouting in *Plantago lanceolata* is based on belowground resource storage (Latzel & Klimesová, 2009; Latzel *et al.*, 2014). Thus, in contrast to pulse response, both clipping tolerance and *R*\* competitive ability are resource-conservative, and therefore both traits should be beneficial in the less nutrient-rich pastures or meadows which make up part of the grassland plots in the Biodiversity Exploratories. However, the two traits were not significantly correlated at the level of individuals or seed families, indicating that they are not physiologically or genetically linked. Another potential explanation for the lack of a family-level relationship could be the inflated genetic variation in the F_1_ generation already explained above (see also Gáspár *et al.*, 2019). However, while F_1_ plants from the same mother may have many different fathers, these most likely come from the same population (Kuiper & Bos, 1992, p. 226), so that population-level differences may have been maintained, and could thus be detected, in our study.

Surprisingly, plant responses to nutrient pulses were negatively correlated to clipping tolerance at the levels of individuals and maternal seed families. Together with the lack of a population differentiation in these traits, this indicates physiological and/or genetic links between them. Again, resource economy appears to be the best explanation. Clipping tolerance is generally thought to be more prevalent in species or genotypes with a more conservative metabolism and more root-, non-structural carbohydrate reserves, whereas a stronger response to a nutrient pulse should requires higher metabolic rate and less storage (Strauss & Agrawal, 1999; Reich, 2014). Thus, there could be a classic resource allocation trade-off between the two traits. The explanation is further supported by the fact that we found the negative correlation mainly in plants from plots with less than one mowing event per year, whereas the relationship tended to disappear at higher mowing frequencies. In the Biodiversity Exploratories, frequent mowing is usually associated with strong fertilisation (Blüthgen *et al.*, 2012). Thus, the resource trade-off disappears when resources become less limiting (Agrawal *et al.*, 2010).

## Acknowledgements

Our work has been funded through the DFG Priority Program 1374 ‘Biodiversity Exploratories’ (DFG grants DU 404/9-1 and BO 3241/2-1). We thank the managers of the three Exploratories, Kirsten Reichel-Jung, Katrin Lorenzen and Martin Gorke, and all former managers, for their work in maintaining the plot and project infrastructure, Christiane Fischer and Jule Mangels for their support through the central office, Andreas Ostrowski and Michael Owonibi for managing the central database, and Markus Fischer, Eduard Linsenmair, Dominik Hessenmöller, Daniel Prati, Ingo Schöning, Francois Buscot, Ernst-Detlef Schulze, Wolfgang Weisser and the late Elisabeth Kalko for their role in setting up the Biodiversity Exploratories project. Fieldwork permits were issued by the responsible state environmental offices of Baden-Württemberg, Thüringen, and Brandenburg (according to §72 BbgNatSchG). We are grateful to Florian Frosch and Jan Helbach for their help during the field campaign, to Christiane Karasch-Wittmann, Eva Schloter, Sabine Silberhorn and Zhiyong Liao for their technical assistance at the University of Tübingen, to Andre Velescu at the Karlsruhe Institute of Technology for soil analyses, and to Niek Scheepens for his support with data analyses.

## Literature Cited

Aarssen LW. 1992. Causes and consequences of variation in competitive ability in plant communities. Journal of Vegetation Science 3:165–174.

Aarssen LW. 2005. On size, fecundity, and fitness in competing plants. In: Reekie EG, Bazzaz FA, eds. Reproductive Allocation in Plants. Academic Press, 215–244.

Aarssen L, Keogh T. 2002. Conundrums of competitive ability in plants: what to measure? Oikos 96:531–542.

Aarssen LW, Turkington R. 1985a. Within-species diversity in natural populations of *Holcus lanatus, Lolium perenne* and *Trifolium repens* from four different-aged pastures. The Journal of Ecology 73:869–886.

Aarssen LW, Turkington R. 1985b. Biotic specialization between neighbouring genotypes in *Lolium perenne* and *Trifolium repens* from a permanent pasture. The Journal of Ecology 73:605–614.

Aarssen LW, Turkington R. 1985c. Competitive relations among species from pastures of different ages. Canadian Journal of Botany 63:2319–2325.

Aarssen LW, Turkington R. 1987. Responses to defoliation in *Holcus lanatus, Lolium perenne*, and *Trifolium repens* from three different-aged pastures. Canadian Journal of Botany 65:1364–1370.

Agrawal AA. 1998. Induced responses to herbivory and increased plant performance. Science 279:1201–1202.

Agrawal AA, Conner JK, Rasmann S. 2010. Tradeoffs and negative correlations in evolutionary ecology. In: M. A. Bell, W. F. Eanes, D. J. Futuyma, and J. S. Levinton, ed. Evolution After Darwin: The First 150 Years. 243–268.

Agrawal AA, Strauss SY, Stout MJ. 1999. Costs of induced responses and tolerance to herbivory in male and female fitness components of wild radish. Evolution 53:1093–1104.

Allan E, Bossdorf O, Dormann CF, Prati D, Gossner MM, Tscharntke T, Blüthgen N, Bellach M, Birkhofer K, Boch S, et al. 2014. Interannual variation in land-use intensity enhances grassland multidiversity. Proceedings of the National Academy of Sciences of the United States of America 111:308–313.

Bergelson J, Crawley MJ. 1992. The effects of grazers on the performance of individuals and populations of scarlet gilia, *Ipomopsis aggregata*. Oecologia 90:435–444.

Bilbrough CJ, Caldwell MM. 1997. Exploitation of springtime ephemeral N pulses by six Great Basin plant species. Ecology 78:231–243.

Blüthgen N, Dormann CF, Prati D, Klaus VH, Kleinebecker T, Hölzel N, Alt F, Boch S, Gockel S, Hemp A, et al. 2012. A quantitative index of land-use intensity in grasslands: Integrating mowing, grazing and fertilization. Basic and Applied Ecology 13:207–220.

Briggs D. 2009. Plant Microevolution and Conservation in Human-influenced Ecosystems. Cambridge University Press.

Coumou D, Rahmstorf S. 2012. A decade of weather extremes. Nature Climate Change 2:491–496.

Davies, M.S. & Snaydon, R.W. 1973. Physiological differences among populations of *Anthoxanthum odoratum* L. on the Park Grass Experiment, Rothamsted. I. Response to calcium. Journal of Applied Ecology 10:33–45.

Davies, M.S. & Snaydon, R.W. 1976. Rapid population differentiation in a mosaic environment. III. Measures of selection pressures. Heredity 36:59–66.

Díaz S, Settele J, Brondízio E, Ngo H, Guèze M, Agard J, Arneth A, Balvanera P, Brauman K, Butchart S, et al. 2019. Summary for policymakers of the global assessment report on biodiversity and ecosystem services of the Intergovernmental Science-Policy Platform on Biodiversity and Ecosystem Services.

Fischer M, Bossdorf O, Gockel S, Hänsel F, Hemp A, Hessenmöller D, Korte G, Nieschulze J, Pfeiffer S, Prati D, et al. 2010. Implementing large-scale and long-term functional biodiversity research: The Biodiversity Exploratories. Basic and Applied Ecology 11:473–485.

Foley JA, Defries R, Asner GP, Barford C, Bonan G, Carpenter SR, Chapin FS, Coe MT, Daily GC, Gibbs HK, et al. 2005. Global consequences of land use. Science 309:570–574.

Gáspár B, Bossdorf O, Durka W. 2019. Structure, stability and ecological significance of natural epigenetic variation: a large-scale survey in *Plantago lanceolata*. The New Phytologist 221:1585–1596.

Gratton C, Denno RF. 2003. Inter-year carryover effects of a nutrient pulse on *Spartina* plants, herbivores, and natural enemies. Ecology 84:2692–2707.

Johnson MTJ. 2011. Evolutionary ecology of plant defenses against herbivores. Functional Ecology 25:305–311.

Juenger T, Bergelson J. 2000. The evolution of compensation to herbivory in scarlet gilia, *Ipomopsis aggregata*: herbivore-imposed natural selection and the quantitative genetics of tolerance. Evolution 54:764–777.

Keddy PA. 1990. Competitive hierarchies and centrifugal organization in plant communities. In: Grace JB, Tilman D, eds. Perspectives on Plant Competition. Academic Press, 265–290.

Kloss L, Fischer M, Durka W. 2011. Land-use effects on genetic structure of a common grassland herb: A matter of scale. Basic and Applied Ecology 12:440–448.

Kuiper PJC, Bos M. 1992. Plantago: A Multidisciplinary Study. Springer.

Latzel V, Klimešová J. 2009. Fitness of resprouters versus seeders in relation to nutrient availability in two *Plantago* species. Acta Oecologica 35:541–547.

Latzel V, Janeček Š, Hájek T, Klimešová J. 2014. Biomass and stored carbohydrate compensation after above-ground biomass removal in a perennial herb: does environmental productivity play a role? Folia Geobotanica 49:17–29.

Lennartsson T, Nilsson P, Tuomi J. 1998. Induction of overcompensation in the field gentian: *Gentianella campestris*. Ecology 79:1061–1072.

Lennartsson T, Tuomi J, Nilsson P. 1997. Evidence for an evolutionary history of overcompensation in the grassland biennial *Gentianella campestris* (Gentianaceae). The American Naturalist 149:1147–1155.

Manning P, Gossner MM, Bossdorf O, Allan E, Zhang Y-Y, Prati D, Blüthgen N, Boch S, Böhm S, Börschig C, et al. 2015. Grassland management intensification weakens the associations among the diversities of multiple plant and animal taxa. Ecology 96:1492–1501.

McNaughton SJ. 1983. Compensatory plant growth as a response to herbivory. Oikos 40:329–336.

Miao SL, Bazzaz FA, Primack RB. 1991. Persistence of maternal nutrient effects in *Plantago major*: the third generation. Ecology 72:1634–1642.

Ostfeld RS, Keesing F. 2000. Pulsed resources and community dynamics of consumers in terrestrial ecosystems. Trends in Ecology & Evolution 15:232–237.

Parepa M, Fischer M, Bossdorf O. 2013. Environmental variability promotes plant invasion. Nature Communications 4:1604.

Pons TL. 1992. Seed germination of Plantago major ssp. major and *Plantago lanceolata*. In: Plantago: A Multidisciplinary Study. Springer, 161–169.

R Development Core Team. 2008. R: A Language and Environment for Statistical Computing.

Reich PB. 2014. The world-wide ‘fast–slow’ plant economics spectrum: a traits manifesto. The Journal of Ecology 102:275–301.

Snaydon RW, Davies MS. 1976. Rapid population differentiation in a mosaic environment. IV. Populations of *Anthoxanthum odoratum* at sharp boundaries. Heredity 37:9–27.

Soliveres S, van der Plas F, Manning P, Prati D, Gossner MM, Renner SC, Alt F, Arndt H, Baumgartner V, Binkenstein J, et al. 2016. Biodiversity at multiple trophic levels is needed for ecosystem multifunctionality. Nature 536:456–459.

Stapledon RG. 1928. Cocksfoot grass (*Dactylis Glomerata* L.): Ecotypes in relation to the biotic factor. The Journal of Ecology 16:71–104.

Strauss SY, Agrawal AA. 1999. The ecology and evolution of plant tolerance to herbivory. Trends in Ecology & Evolution 14:179–185.

Tilman D. 1985. The resource-ratio hypothesis of plant succession. The American Naturalist 125:827–852.

Tokeshi M. 2009. Species Coexistence: Ecological and Evolutionary Perspectives. John Wiley & Sons.

Völler E, Auge H, Bossdorf O, Prati D. 2013. Land use causes genetic differentiation of life-history traits in Bromus hordeaceus. Global Change Biology 19:892–899.

Völler E, Bossdorf O, Prati D, Auge H. 2017. Evolutionary responses to land use in eight common grassland plants. The Journal of Ecology 105:1290–1297.

Warwick SI, Briggs D. 1978. The Genecology of Lawn Weeds. I. Population differentiation in *Poa annua* L. in a mosaic environment of bowling green lawns and flower beds. The New Phytologist 81:711–723.

Warwick SI, Briggs D. 1979. The Genecology of Lawn Weeds. III. Cultivation experiments with *Achillea millefolium* L., *Bellis perennis* L., *Plantago lanceolata* L., *Plantago major* L. and *Prunella vulgaris* L. collected from lawns and contrasting grassland habitats. The New Phytologist 83:509–536.

Weiner J. 1988. The influence of competition on plant reproduction. In: Lovett Doust J, Lovett Doust L, eds. Plant reproductive ecology: patterns and strategies. Oxford University Press, 228–245.

Wilson JB, Spijkerman E, Huisman J. 2007. Is there really insufficient support for Tilman’s *R\** concept? A comment on Miller et al. The American Naturalist 169:700–706.

Yang LH, Bastow JL, Spence KO, Wright AN. 2008. What can we learn from resource pulses? Ecology 89:621–634.

Zuur AF, Ieno EN, Walker NJ, Saveliev AA, Smith GM. 2009. Mixed effects models and extensions in ecology with R. Springer.

